# An omega glutathione S-transferase in *Apis mellifera* contributes to chemical adaptation through pesticide sequestration and antioxidant defense

**DOI:** 10.64898/2026.03.03.709375

**Authors:** Sonu Koirala B K, Timothy W. Moural, Gaurab Bhattarai, Ngoc T Phan, Edwin G Rajotte, David J Biddinger, Fang Zhu

## Abstract

The European honey bee (*Apis mellifera* L.) is a key agricultural pollinator frequently exposed to pesticide residues, yet the molecular basis of its chemical adaptation, particularly glutathione S-transferases (GSTs) involved in xenobiotic detoxification, remain incompletely understood. In this study, AmGSTO1 was structurally and functionally characterized to evaluate its role in agrochemical interaction and protection against oxidative stress. The crystal structure of AmGSTO1 in complex with glutathione revealed its 3D architecture and key active-site residues were identified by structural analysis and site-directed mutagenesis. Fluorescence binding assays demonstrated measurable affinity for multiple agrochemicals, including TCP, fenoprop, 2,4-D, tetramethrin, nicotine, and 3-phenoxybenzaldehyde. However, HPLC analysis showed no detectable substrate depletion, suggesting ligand binding to AmGSTO1 without catalytic turnover. AmGSTO1 exhibited antioxidant activity toward cumene hydroperoxide, hydrogen peroxide, and paraquat, as well as dehydroascorbate reductase activity. These findings indicate that AmGSTO1 may contribute to agrochemical tolerance through ligand sequestration and redox protection mechanisms.

## INTRODUCTION

The European honey bee (*Apis mellifera* L.) has made significant ecological and economic contributions to human civilization through both their hive products and pollination services ^1-2^. However, in recent decades, several potential threats that directly impact honey bee health and their ecosystems have been identified, including pesticides, climate change, nutrient deficits, pests, pathogens, and their interactions ^3-8^. Ultimately, these threats lead to reduced agricultural productivity ^9^. Among different stressors, there is substantial evidence that honey bees are exposed to a variety of agrochemicals, both within their hive surroundings and through pollen and nectar that they consume ^10^. Previous study indicated that, on average, North American bee colonies are contaminated with 6.5 different pesticides ^11^. Depending on the route of exposure, mode of action, and dose, pesticides may induce acute toxicity, chronic toxicity, or no toxicity in bees ^12-14^. Additionally, bees that consume plant allelochemicals in their diet may enhance their ability to detoxify pesticides, thereby improving their resilience to chemical stress ^15-16^. Therefore, it is important to understand the molecular mechanisms by which bees adapt to environmental toxins ^14^. Compared to other insects, honey bees have a lower number of detoxification enzymes, which may lead to greater sensitivity toward pesticides and other toxins in the environment ^17^. Nevertheless, they are capable of metabolizing a variety of insecticides through enzymatic mechanisms, although the complete spectrum of chemicals they can process is still poorly understood ^18^.

Similar to other insects, the honey bee genome is comprised of numerous detoxification genes, including glutathione S-transferases (GSTs), a group of enzymes that play an important role in the detoxification of numerous endogenous and exogenous compounds ^17, 19-20^. In general, GSTs catalyze the conjugation of a wide range of electrophilic substances and toxic compounds to glutathione. This process facilities metabolism, detoxification, and elimination of numerous pesticides and plant toxins ^21-22^. In addition to their roles in direct metabolism, GSTs may participate in passive non-catalytic binding of substances (i.e. sequestration) ^22-23^. Moreover, they exhibit peroxidase activity, helping to reduce oxidative stress caused by biotic and abiotic stressors ^20, 24^. Insect cytosolic GSTs have been characterized and classified into six subclasses: delta, epsilon, omega, sigma, theta, and zeta ^25-28^. The honey bee genome contains 10 cytosolic GST genes, among which AmGSTO1 has been identified as an omega-class GST ^17^. Omega GSTs are cysteine-type GSTs, with a cysteine residue that interacts with GSH by forming a disulfide bond ^29^. In contrast, other cytosolic GSTs use tyrosine and serine for GSH stabilization and as the key residues in the active site necessary for metabolism of toxic chemical compounds ^26, 30^. Furthermore, Omega GSTs have been linked to red eye development via the metabolism of eye pigments in *Drosophila melanogaster* Meigen, demonstrating that omega-class GSTs function in biosynthetic pathways ^31^. In humans, the omega-class GSTs may have a broader role in ion channel modulation beyond their well-established functions in metabolism and detoxification ^32^. In pollinators, studies on omega-class GSTs remain limited, with little information available on their structural characteristics, substrate specificity, or overall role in xenobiotic adaptation.

In this study, we investigated the structure and function of AmGSTO1 to clarify its role in xenobiotic adaptation in honey bees. We characterized its tissue-specific expression, amd resolved its three-dimensional co-crystal structure with GSH by X-ray crystallography. Using structural analysis and site-directed mutagenesis, we identified its key active-site residues. Moreover, its antioxidative properties against cumene hydroperoxide, hydrogen peroxide, and paraquat were assessed with cellular assays, including disc diffusion and bacterial survival assays. We further quantified its binding affinities to a broad panel of pesticides, metabolites, and plant allelochemicals, and evaluated its catalytic activity toward selected agrochemicals using high-performance liquid chromatography (HPLC). Together, these analyses offer a comprehensive view of an omega-class GST in honey bees and highlight its potential role in pesticide sequestration and response to oxidative stress.

## MATERIALS AND METHODS

### Phylogenetic Analysis of AmGSTO1 and Other Insect GSTs

The full coding sequence of the AmGSTO1 gene was cloned using sequencing information from the NCBI database (XP_006569695.1) and primers listed in Table S1. PCR-amplified cDNA was purified, T4-treated, and ligated into a HpaI-digested pET-9BC vector, transformed into DH5α cells, and positive colonies were verified by plasmid extraction and sequencing (Functional Biosciences, WI, USA). To classify AmGSTO1, a phylogenetic tree was constructed using 157 GST amino acid sequences (Table S2) from 5 insect species (characterized and predicted) retrieved from NCBI: *A. mellifera, D. melanogaster, Tribolium castaneum* (Herbst), *Apis cerena cerena* F., and *Bombus impatiens* Cressson. Multiple protein sequence alignment was performed using ClustalW ^33^ in MEGA X ^34^ with default parameters (gap open penalty: 10, gap extension penalty: 0.2). The maximum likelihood ^35^ unrooted phylogenetic tree was inferred using RaxML 8.2.12 ^36^ using PROTGAMMABLOSUM62 model which uses the BLOSUM62 substitution matrix and incorporates a gamma distribution to account for varying rates of evolution across sites, with bootstrap analysis performed using 500 replicates to infer the consensus tree.

### RNA Extraction, cDNA Synthesis, and qRT-PCR

Nurse and forager bees were collected from three hives spaced 60 m apart at the Penn State Wiley Apiary in University Park, PA, which were managed using standard best practices (https://extension.psu.edu/best-management-practices-for-bee-health; accessed on 20 December, 2025). Nurse bees were taken from the brood nest, and foragers were collected by shaking honey frames ^37^. Then the bees were flash frozen in liquid nitrogen and stored at - 80°C. Frozen nurse and forager bees were dissected in phosphate-buffered saline solution (0.01M, pH 7.4), and the head, fat body, Malpighian tubule, midgut, legs, and muscle were collected. Total RNA was extracted using Invitrogen Trizol reagent (Invitrogen, Carlsbad, CA, USA) following the manufacturer’s protocol. For whole-body analysis, a biological replicate containing four nurse or forager bees, which were ground separately in liquid nitrogen prior to RNA extraction. For tissue analysis, each replicate comprised pooled tissues from 5–15 bees. Three biological replications were used for each experiment. RNA quality and purity were assessed with a NanoDrop™ One spectrophotometer (Thermo Fisher Scientific Inc., Waltham, MA, USA), using a A260/280 ratio of 1.8-2.0 as the quality criterion. cDNA synthesis was performed with M-MLV Reverse transcriptase (Thermo Fisher Scientific, Waltham, MA), then cDNA was used as a template for qRT-PCR reaction. Each 10 µL reaction contained 1 μL cDNA, 5 μL FastStart SYBR Green Master (Roche Diagnostics, Indianapolis, IN USA), 0.4 μL qRT-PCR primers (Table S1), and 3.6 μL ddH_2_O. Reactions were run on a Bio-Rad CFX Connect™ Real-Time PCR System (Bio-Rad, CA, USA). The thermal cycling conditions were 95°C for 10 min., followed by 39 cycles of 95°C for 10s and 55°C for 30s. The most stable housekeeping gene, GADPH (Glyceraldehyde-3-phosphate dehydrogenase) (XM393605), was used for normalization of expression of *AmGSTO1* using the 2-ΔΔCT method ^38-39^. The overall difference in the level of expression among the tissues was analyzed by one-way ANOVA with the Tukey HSD test for multiple comparisons in R (Version 4.1.0). The difference between nurse and forager whole body expression was compared by Student’s t-test.

### Recombinant AmGSTO1 Expression and Purification

Honey bee AmGSTO1 (XP_006569695.1) with a 6xHis tag was ligated into the circular plasmid vector (Addgene #48285 plasmid, pET-9BC) via ligation independent cloning. Then the newly constructed plasmid was transferred into the Rosetta™ II (DE3) pLysS BL21 *Escherichia coli* expression strain. For protein expression, *E. coli* containing pET9BC-AmGSTO1 was incubated at 37 ℃ overnight at 250 rpm in 100 ml terrific broth media with antibiotics ampicillin (200 µg/ml) and chloramphenicol (CAM) (30 µg/ml) ^40^. Then the culture was inoculated in 2 L of TB media which underwent incubation at 37 ℃ until its optical density (OD) at 600 nm reached between 0.4 and 0.6. AmGSTO1 expression was induced by the addition of 0.5 mM IPTG and incubated for another 24 h at 20 ℃ by shaking at 250 rpm. Cell pellets were harvested by centrifuging at 4000 rpm, 4 ℃ and stored at -20 ºC. AmGSTO1 was purified as described previously with some modifications ^39^. In brief, frozen cell pellets were resuspended in lysis buffer containing 25 mM NaPi, 500 mM NaCl, 3.3 mM NaN_3_, and 20 mM imidazole at pH of 7.6 mixed with 1 mM PMSF, 1mM DTT, and a protease inhibitor tablet (Thermo Scientific^™^) at pH 7, then lysed by sonication (Branson Digital Sonifier SFX 150). The lysate was centrifuged at 18,500 rpm, 4 °C to separate water-soluble protein and insoluble materials. The soluble protein was injected into an NGC Medium-Pressure Liquid Chromatography (MPLC) System (Bio-Rad Laboratories, Hercules, CA, USA) connected to a 5 mL Co-NTA column. Then 6xHis tag fused AmGSTO1 was eluted with 20 mM NaPi, 300 mM NaCl, 3.3 mM NaN_3_, and 250 mM imidazole at pH of 7.6. The resulting protein was subjected to a 100-fold buffer exchange, using a centrifugal concentrator with a molecular weight cutoff (MWCO) of 10 kDa, into a buffer solution containing 5 mM NaPi, 5 mM HEPES, and 5mM DTT at pH of 7.6. Buffer exchanged protein was further purified by a Hydroxyapatite column (HA, ceramic hydroxyapatite type I, BIO-RAD) connected to an NGC MPLC system. Then the gradient ranging from 5 mM NaPi, 5 mM HEPES, and 5 mM DTT at pH of 7.6 to 500 mM NaPi and 3.3mM NaN3 at pH of 7.6 was used to wash and elute the recombinant protein. The protein was buffer exchanged with a solution containing 20 mM Tris at pH 7.6 using a centrifugal concentrator (10 kDa MWCO) and then introduced to an Enric™ Q 10 100 mm high resolution ion exchange column (BIO-RAD). The gradient ranging from 20 mM Tris at pH of 7.6 with 2 M DTT, to 20 mM Tris, 1 M NaCl, and 2 mM DTT at pH 7.6 was used to wash and elute the protein. Then, the final purification step was performed with a Cytiva® HiPrep Sephacryl S-200 HR size exclusion column with 20 mM HEPES, 150 mM NaCl, 1 mM EDTA, and 20 mM GSH at pH 7.6. Protein obtained from various columns was followed by SDS-PAGE and concentration determination using a NanoDrop™ One (Invitrogen) spectrophotometer to validate the quality and quantity of protein (Fig. S1). The protein was flash-frozen in liquid nitrogen and stored at -80 ºC for future use.

### X-ray Crystallography of AmGSTO1-GSH complex and molecular docking with CDNB

AmGSTO1 was crystalized by hanging drop vapor diffusion at 18 °C. Protein (5 mg/ml) with 20 mM GSH was mixed 1:1 with a reservoir solution (0.2 M ammonium acetate and 20% PEG 3,350) in VDX plates. The co-crystals were placed into cryoprotection solution (0.2M ammonium acetate, 20% PEG 3,350, 30% glycerol) and x-ray diffraction data were collected at the Advanced Photon Source Structural Biology Centers beamline 19-ID. Software such as Collaborative Computational Project Number 4 (CCP4) with DIALS, and xia2 software was used for processing diffraction data ^41-45^. Phaser implemented in Phenix was used for phasing using an AlphaFold2 model as a search model for molecular replacement ^46-48^. Model building and refinement was performed using Phenix and Coot ^49-50^. Structural analysis and figures were generated following previously published methods ^39^. For molecular docking, the structure of 1–chloro-2, 4-dinitrobenzene (CDNB) was downloaded from the PubChem® database in SDF format ^51^. Ligands were docked using AutoDock Vina in UCSF Chimera, with the prepared chain A structure (GSH-bound) used as the receptor ^52-53^. The substrate-binding pocket was used as the docking site. Nine poses were generated for CDNB, and one representative pose was selected based on the AutoDock Vina docking score, with more negative scores indicating stronger binding affinity. Figures with docked ligands were prepared in ChimeraX ^54^.

### Enzymatic Assay

To understand the enzymatic property and its substrate range, the kinetics analysis of purified recombinant AmGSTO1 was conducted using various substrates, including CDNB, reduced glutathione (GSH), p-nitrophenyl acetate (PNA) and dehydroascorbic acid (DHA) ^55-56^. Stock solutions (100 mM) CDNB and PNA were prepared in ethanol, DHA in DMSO, and GSH in a KPi buffer. CDNB-conjugating activity was assayed by varying CDNB concentrations (0.125 -1.25 mM) at constant 1mM GSH. PNA activity was measured with substrates ranging from 0.0625 to 1.5 mM (GSH fixed at 0.5 mM). GSH assayed using 0.031 - 0.75 mM GSH and CDNB fixed at 1 mM. DHA reductase activity was measured using DHA at 1 mM and 1 mM GSH. The concentration of AmGSTO1 was 0.2mg/mL for CDNB and GSH, and 0.8 mg/ml for PNA and DHA, in 100 mM KPi buffer (pH 8). Reactions were carried out in 96-well UVstar® microplates (Greiner Bio-One). The assays were conducted in triplicate with a reaction volume of 200 μL. Absorbance at 340 nm (CDNB), 405 nm (PNA), and 265 nm (DHA) ^55-56^ was monitored for 1 minute 10 seconds using a Spark® multi-mode plate reader (TECAN, Mannedorf, Switzerland). Enzyme-free controls were included to correct for non-enzymatic reaction. Kinetic parameters were calculated using published extinction coefficients with 96-well pathlength correction and analyzed in GraphPad Prism ^21, 55, 57^.

### Site-Directed Mutagenesis

To examine the contribution of active-site residues to enzyme function, Cys28, Tyr30, Glu81, Ser82, Phe127, and Trp174 of AmGSTO1 were mutated to Ala. PCR primers were designed using NEBaseChanger (New England Biolabs; https://nebasechangerv1.neb.com/) to introduce specific nucleotide substitutions or modifications for AmGSTO1 (Table S1). Amino acid substitution variants of AmGSTO1 were generated utilizing the Q5® Site-Directed Mutagenesis Kit (New England Biolabs®, MA, USA). Wild-type recombinant AmGSTO1-pET9BC was used as the template for generating mutants, and full-length mutant constructs were verified with DNA sequencing. Mutant AmGSTO1-pET9BC was introduced into Rosetta™ II (DE3) pLysS cells for expression, and proteins were purified using the same procedure as wild-type AmGSTO1. Purified mutant AmGSTO1 protein was then subjected to GSH-CDNB kinetic assay, as described in previous section using mutant AmGSTO1 at a concentration of 0.8 mg/mL.

### Fluorescence Binding Assay

Fluorescence binding assay was performed to examine H-site interactions of GSTs Using 8-Anilinonaphthalene-1-sulfonic acid (ANS) as the probe (excitation: 380/20 nm emission: 485/20 nm bandwidth). ANS saturation binding assay was performed with AmGSTO1 to determine the maximum binding capacity (B_max_) and equilibrium dissociation constant (*K*_d_). 2 μM AmGSTO1 was titrated with 0 - 225 μM ANS, and curve fitted in GraphPad Prism 9.5.1. ANS displacement assay assessed competition by other substrates. For displacement assay, AmGSTO1 (2 μM) and ANS (50 μM) were incubated with competitor ligands (0 - 4 mM) in 20 mM potassium phosphate buffer containing 150 mM NaCl, 1 mM EDTA, and 1 mM Tris(2-carboxyethyl) phosphine (TCEP) (pH 6) in 200 μL reactions in 96-well black plates. Measurements were performed using a Tecan Spark® multi-mode plate reader, with CDNB and ethacrynic acid (EA) as positive controls and GSH as a negative control for ANS displacement. Multiple herbicides (atrazine, clopyralid, 2,4-D, dicamba, fenoprop, metolachlor, paraquat, picloram, 3,5,6-trichloro-2-pyridinol [TCP], triclopyr), fungicides (chlorothalonil, propiconazole, pyraclostrobin, myclobutanil), insecticides and their metabolites (acephate, amitraz, bifenthrin, carbaryl, 6-chloronicotinic acid, chloropyrifos, coumaphos, deltamethrin, diazinon, dinotefuran, fenvalerate, imidacloprid, malathion, metamidophos, permethrin, 3-phenoxybenzaldehyde, tetramethrin), and plant allelochemicals (caffeine, cotinine, nicotine, p-coumaric acid, phenethyl isothiocyanate, phenylacetaldehyde, propyl isothiocyanate) were used as competitors in ANS displacement assays. Each reaction was performed in triplicate. Finally, IC_50_ values for competitive ligands were calculated using GraphPad Prism 9.5.1, and dissociation constants (*K*_i_) were determined using the equation *K*_i_ =[IC_50_]/(1 + [ANS]/*K*_d_^ANS^) ^58^.

### Metabolism Assay by HPLC MS/MS

To assess the catalytic property of AmGSTO1, the enzymatic assay for the HPLC study was conducted in a reaction containing 0.02 M ammonium acetate, 4 mM GSH, 0.2 mM of each pesticide (dissolved in methanol), and 2 μM of the enzyme in a total volume of 500 µL. Following a one-hour incubation, the reaction was halted by the addition of 500 µL of methanol. Subsequently, the mixture underwent separation 3K centrifugal filter units. The filtrate was then introduced into the high sensitivity Vanquish Flex UHPLC system for LC-MS and LC-MS/MS analyses located in the Huck Core Facility (Penn State University). A non-enzymatic reaction control was carried out comprising same amount of heat-inactivated enzyme and all other reactants used in treatment groups. Each reaction, for both controls and treatments, was performed in triplicate (n = 3). The chemicals included fenoprop, TCP, and 2,4-D. Detection of chemical constituents was achieved through a UV detector, while the obtained mass spectra were cross-referenced with the NIST Mass Spectral Database version 2.0 (NIST, Gaithersburg, MD) to elucidate the chemical composition within the mixture.

### Disc Diffusion Assay

To access the antioxidative properties of AmGSTO1, a disc diffusion assay was performed with a method modified from Burmeister’s protocol ^59^. Transformed cells with recombinant AmGSTO1-pET-9BC plasmid was used as the treatment group, and the empty plasmid (pET-9BC) vector served as control. For control and treatment transformed Rosetta™ II (DE3) pLysS cells were grown in LB with ampicillin (200 µg/ml) and chloramphenicol (30 µg/ml) at 37 °C, 250 rpm. When culture reached OD_600 nm_ = 0.6, cells were induced with 1 mM of IPTG. Next, the cultures were incubated at 37 °C for 7 hours. Approximately 5 × 10^8^ cells were plated on LB agar with ampicillin (200 µg/ml) and chloramphenicol (30 µg/ml). Then plates were incubated at 37 °C for 1h to dry the plates. After that, filter discs (6-mm diameter) soaked with different concentrations (0 mM, 50 mM, 100 mM, 150 mM, and 200 mM) of cumene hydroperoxide (CHP) and hydrogen peroxide (H_2_O_2_) were placed on the plates separately. Then, after 12 hours of incubation at 37 °C, the halo diameter for control and treatment plates were recorded. The differences in the diameter of the halo were analyzed by one-way ANOVA with the Tukey HSD test for multiple comparisons in R (Version 4.1.0).4.

### Bacterial Survival Assay

To assess the role of AmGSTO1 against oxidative stress, a bacterial survival assay was performed following Dong’s protocol ^60^. H_2_O_2_ and paraquat were dissolved in water. *E. coli* carrying empty plasmid (pET-9BC) vector was used as the control and cells with recombinant AmGSTO1-pET-9BC were used as the treatment. Treatment and control cells (5 mL) were cultured separately in 2YT media containing ampicillin (200 µg/mL) and CAM (30 µg/mL) at 37 °C, 180 rpm. When cultures reached OD_600 nm_ = 0.3 nm, 1 mM IPTG and 0.5 mM H_2_O_2_ or paraquat were added to the control or treatment cells. After 1 h of induction with oxidative agents at 37 °C, 180 rpm, the OD_600 nm_ was measured hourly. After 7-8 hours of incubation, bacterial suspensions were diluted 10,000-fold in autoclaved water, and 50 μL of the bacteria was spread on LB agar plates containing ampicillin (200 µg/mL) and CAM (30 µg/mL). Colony-forming units (CFUs) were recorded after overnight incubation at 37 °C. All assays were repeated with three biological replicates, and differences between control and treatment groups were analyzed using Student t-test.

## RESULTS

### Phylogenetic Relationships of AmGSTO1 with Other Insect GSTs

The phylogenetic analysis of AmGSTO1 (accession number XM_006569632.2) was performed to investigate the evolutionary relationships of AmGSTO1 with other insect GST genes (Table S2) and to establish orthology with other insect GSTs. The AmGSTO1 sequence has an open reading frame of 726 bp, encoding a 242 amino acid protein. Phylogenetic analysis grouped GST enzymes by class. AmGSTO1 clustered into the omega clade alongside an omega class GST from *A. cerena cerena* (AccGSTO1) (XP_016918436.1) and its predicted isoform (AGX26610.1), along with one omega GST from *Bombus impatiens* (XP_012237875.1) (Fig. 1).

**Fig. 1.**
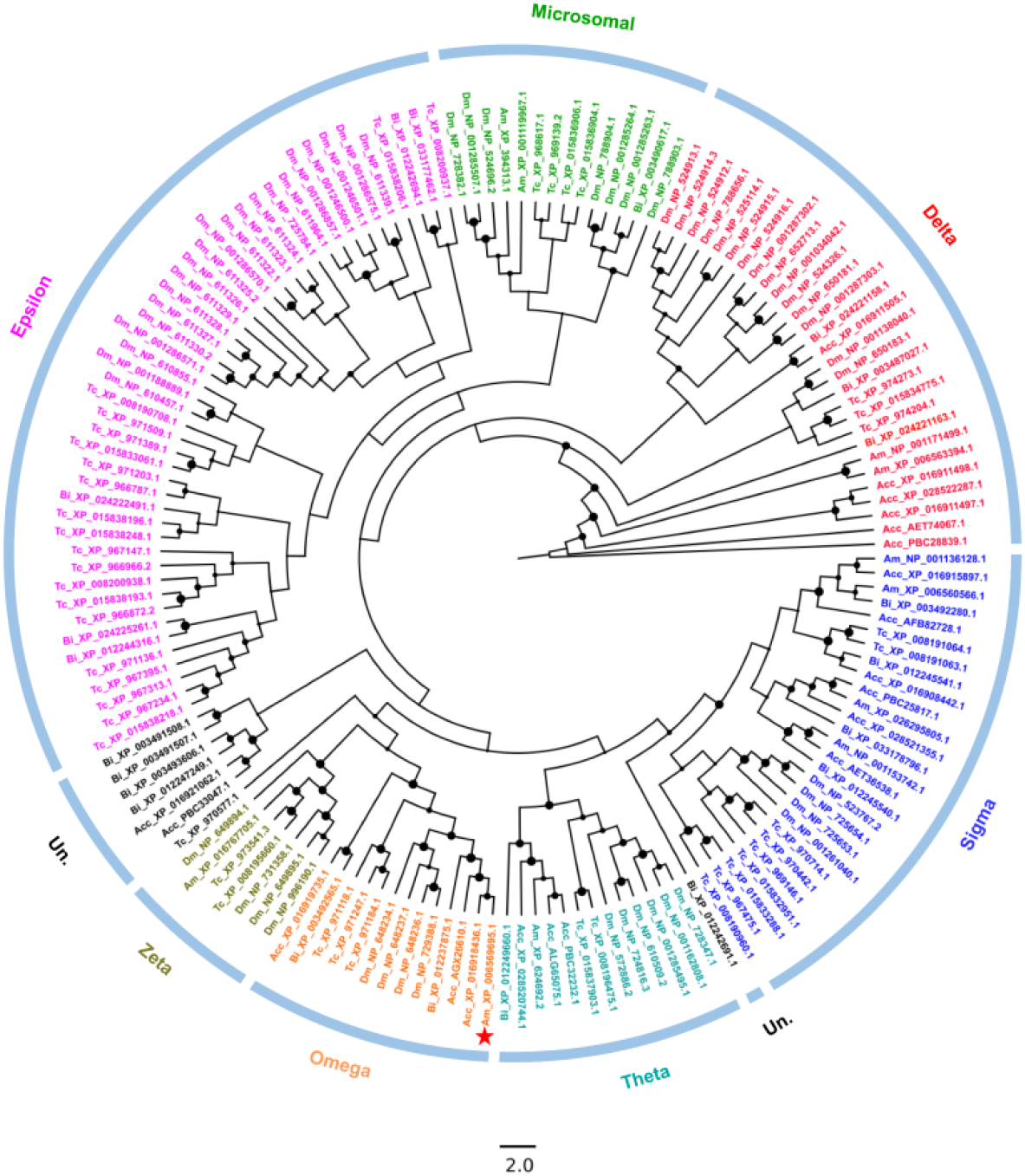
Unrooted maximum likelihood tree of 157 GST proteins in five different insect species: *Drosophila melanogaster* ^45^, *Tribolium castaneum* (Tc), *Apis mellifera* (Am), *Apis cerana cerana* (Acc), and *Bombus impatiens* (Bi). The color of terminal nodes represents cytosolic GST from various classes. GST proteins in black indicate uncharacterized ones. The size of solid circles at each node represents the bootstrap support value. Accession indicated with a star symbol indicates omega-class GST (AmGSTO1) in honey bees.

### Gene Expression Patterns of *AmGSTO1* in Nurse and Forager Bees

qRT-PCR analysis was carried out to examine the temporal and spatial expression patterns of *AmGSTO1* in nurse and forager bees. The relative expression of *AmGSTO1* in forager bees was significantly higher than nurse bees (*p*<0.001) (Fig. 2A). In terms of spatial expression, we found a significant difference in *AmGSTO1* expression among different tissues in both nurse (*p*<0.001; df = 5; F = 113.52) and forager (*p*<0.001; df = 7; F = 30.96) bees. Specifically, in nurse bees, the mean relative expression of *AmGSTO1* was significantly higher in the fat body, followed by the Malpighian tubules, head, midgut, legs, and muscle (Fig. 2B). In forager bees, the expression of *AmGSTO1* was significantly higher in fat body followed by hind legs, Malpighian tubules, fore legs, and head, with comparatively lower expression in the middle legs, midgut, and muscle (Fig. 2C).

**Fig. 2.**
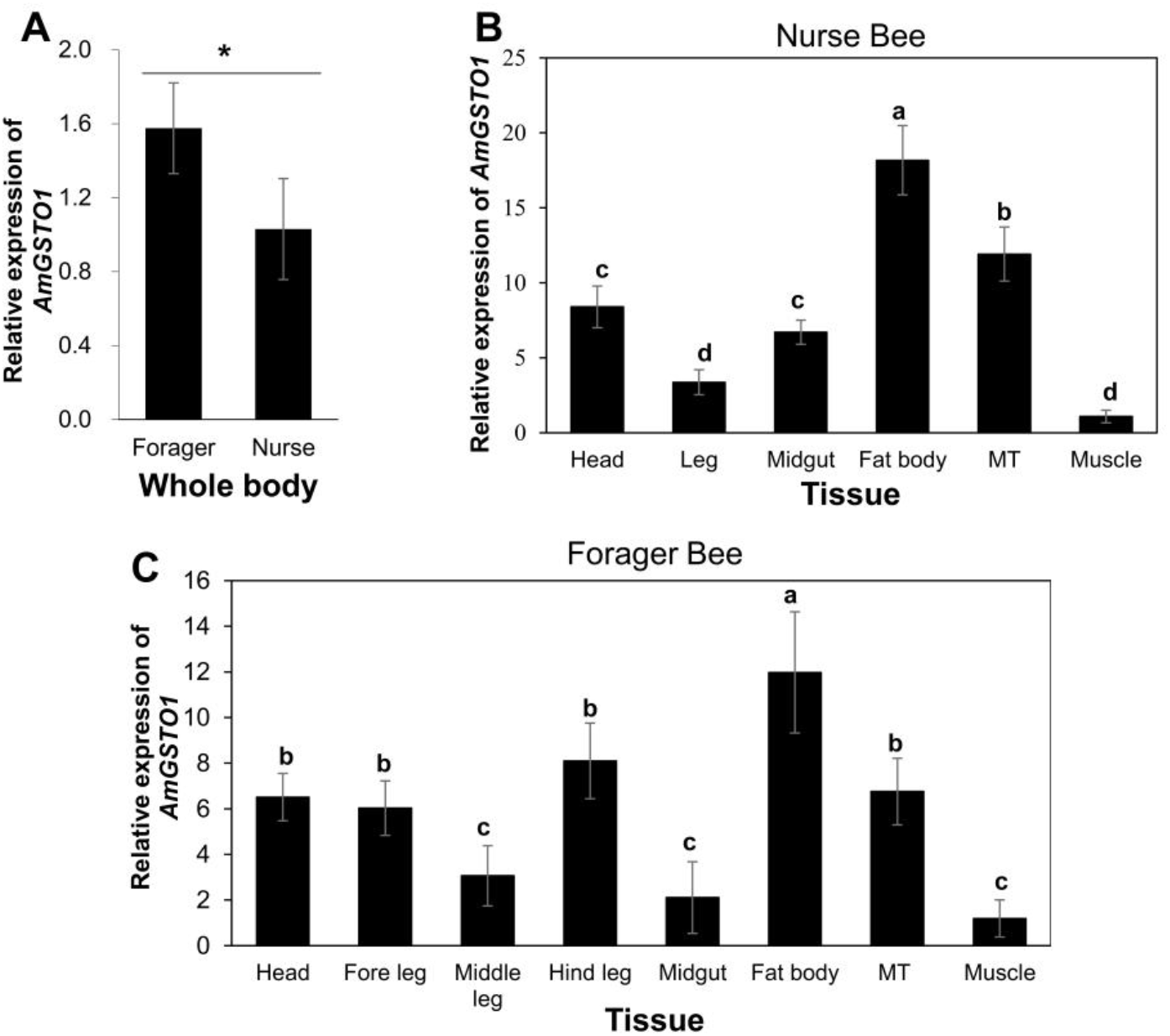
Gene expression patterns of *AmGSTO1* in nurse and forager bees. (A) Whole body gene expression pattern. Spatial expression profile in nurse bees (B) and forager bees (C). Different letters indicate a significant difference in gene expression at p < 0.001, according to one-way ANOVA with the Tukey HSD test. * Indicates a significant difference of gene expression between nurse and forager bees by Student’s t-test analysis. Note: MT-Malpighian tubules.

### X-ray Crystal Structure of AmGSTO1 and Its Co-crystal with GSH

A protein bank database (PDB) search with the AmGTSO1 sequence revealed the highest sequence match with insects was *Bombyx mori* omega-class GST (3WD6) with a sequence identity of 35% ^61^. The asymmetric unit contained four molecules of AmGSTO1 each complexed with one GSH molecule. AmGSTO1 crystallized in space group P 2_1_, where the unit cell dimensions were a = 64.14 (Å), b = 78.39 (Å), c = 107.97 (Å), with angles α = 90.00°, β = 106.70°, and γ = 90.00°. The structure was refined out to 2.05 (Å), with R_work_ and R_free_ values of 22.10% and 25.78%, respectively (Table 1). An analysis of the diffraction data indicated that AmGSTO1 forms a dimeric protein structure. The dimeric protein comprises two monomers designated as subunit 1 and subunit 2 (Fig. 3). The AmGSTO1 monomer comprises N-terminal and C-terminal domains connected by a 10-amino acid linker. The structure exhibits a typical thioredoxin-like fold, with an N-terminal comprising β_1_α_1_β_2_α_2_β_3_β_4_α_3_ motifs and C-terminal helical domain. N-terminal domain was ordered starting with β_1_ (residues I20-S24), followed by α_1_ (residues P29-A40), β_2_ (residues H45-Y49), α_2_ (residues D57-K62), β_3_ (residues C70-E72), β_4_ (residues I78–Y80), and α_3_ (residues S82-T92). The C-terminal domain commenced with α_4_ (residues P103-I128), followed by α_5_ (residues Q134-R154), α_6_ (residues M166-R185) with a bulge at R177, α_7_ (residues D187-F189), α_8_ (residues K197-E208) with a bulge at N209, and α_9_ (residues P210-N215), and α_10_ (residues T219-R230) (Fig. 3). Each subunit contains two ligand-binding sites: a conserved G-site in Domain I for binding GSH and an H-site in Domain II for hydrophobic substrates.

**Table 1.**
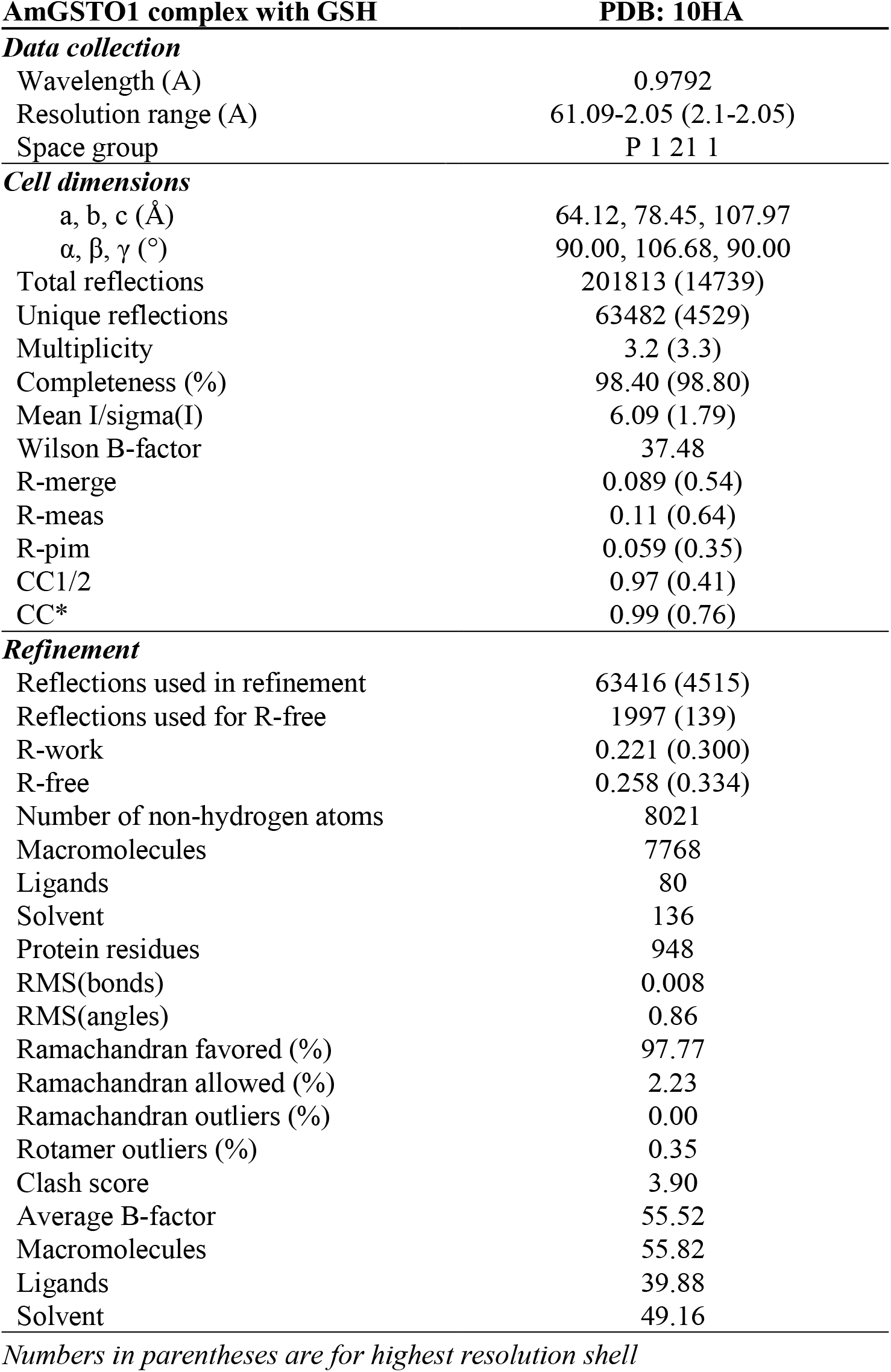
Data collection and refinement for AmGSTO1 and GSH.

| **AmGSTO1 complex with GSH** | **PDB: 10HA** |
| --- | --- |
| ***Data collection*** |  |
| Wavelength (Å) | 0.9792 |
| Resolution range (Å) | 61.09-2.05 (2.1-2.05) |
| Space group | P 1 21 1 |
| ***Cell dimensions*** |  |
| a, b, c (Å) | 64.12, 78.45, 107.97 |
| $\alpha, \beta, \gamma$ (°) | 90.00, 106.68, 90.00 |
| Total reflections | 201813 (14739) |
| Unique reflections | 63482 (4529) |
| Multiplicity | 3.2 (3.3) |
| Completeness (%) | 98.40 (98.80) |
| Mean I/sigma(I) | 6.09 (1.79) |
| Wilson B-factor | 37.48 |
| R-merge | 0.089 (0.54) |
| R-meas | 0.11 (0.64) |
| R-pim | 0.059 (0.35) |
| CC1/2 | 0.97 (0.41) |
| CC* | 0.99 (0.76) |
| ***Refinement*** |  |
| Reflections used in refinement | 63416 (4515) |
| Reflections used for R-free | 1997 (139) |
| R-work | 0.221 (0.300) |
| R-free | 0.258 (0.334) |
| Number of non-hydrogen atoms | 8021 |
| Macromolecules | 7768 |
| Ligands | 80 |
| Solvent | 136 |
| Protein residues | 948 |
| RMS(bonds) | 0.008 |
| RMS(angles) | 0.86 |
| Ramachandran favored (%) | 97.77 |
| Ramachandran allowed (%) | 2.23 |
| Ramachandran outliers (%) | 0.00 |
| Rotamer outliers (%) | 0.35 |
| Clash score | 3.90 |
| Average B-factor | 55.52 |
| Macromolecules | 55.82 |
| Ligands | 39.88 |
| Solvent | 49.16 |

*Numbers in parentheses are for highest resolution shell*

**Fig. 3.**
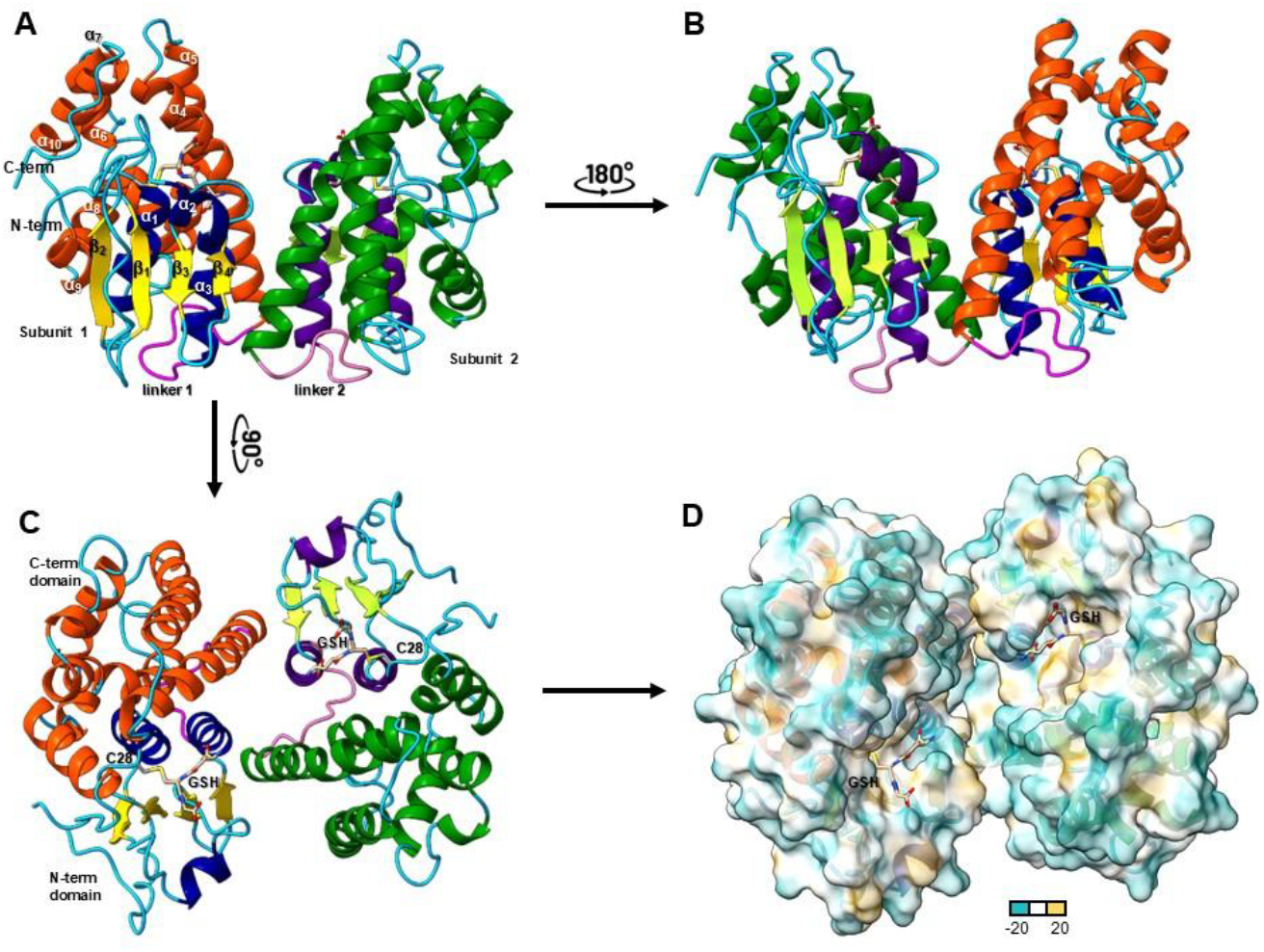
Global structure of AmGSTO1 shown as a dimer and represented in ribbon diagram and surface overlayed on a ribbon diagram. (A) Dimeric ribbon diagram of AmGSTO1 showing Subunit 1 (monomer 1) and Subunit 2 (monomer 2). (B) The ribbon diagram of the dimeric AmGSTO1 structure from A has been rotated 180° showing the two-fold symmetry of the dimer. (C) The ribbon diagram of the dimeric AmGSTO1 structure from A rotated 90°. (D) The dimer ribbon diagram overlayed with the lipophilic surface. The lipophilic representation scale is shown with -20 being the least lipophilic and 20 being the most lipophilic. Figures were generated with UCSF Chimera X v:1.5.

### Active site determination

The AmGSTO1-GSH co-crystal structure revealed a hydrophilic G-site in the N-terminal domain interacting with GSH, adjacent to a lipophilic pocket for hydrophobic substrate binding (Fig. 3D). Structural analysis indicates that the glutathionyl moiety of the ligand forms hydrogen bonds with residues Lys67, Val68, Glu81, and Ser82 with interaction distances ranging from 2.6 to 3.4 Å (Fig. 4A). The sulfur atom from glutathione’s cysteine residue formed a mixed disulfide with the active-site cysteine-28 (Cys28) of AmGSTO1 (Fig. 4A,C). Residues Glu81 and Ser82 participate in binding with the γ-glutamyl moiety of GSH. The Glu81 OE1 atom lies 3.01 Å from the GSH γ-glutamyl N1 atom, while the Ser82 OG and N atoms are positioned 2.92 Å from the γ-glutamyl O12 atom, respectively. Additional hydrogen bond interactions were between the glycyl O32 of GSH and Lys55 NZ of AmGSTO1 (2.74 Å), along with a hydrogen bond between the GSH cysteine carbonyl oxygen O2 and Val68 of AmGSTO1 (3.02 Å). The surrounding GSH-binding pocket is formed by Tyr30, Leu52, Lys55, Gly66, and Pro69. Among the generated docking poses, the CDNB with the lowest binding energy (–6.09 kcal/mol) was selected, with C3 located 4.68 Å from the glutathione sulfur atom. The H-site is located adjacent to the G-site, composed of Pro29, Tyr30, Cys124, Phe127, Ile128, Trp174, Arg177, Try225, Met226, and Arg229 (Fig. 4B). Conserved omega class GST residues in the G-site were found to be, Cys28, Pro29, Arg33, and Leu38 (Fig. 3S).

**Fig. 4.**
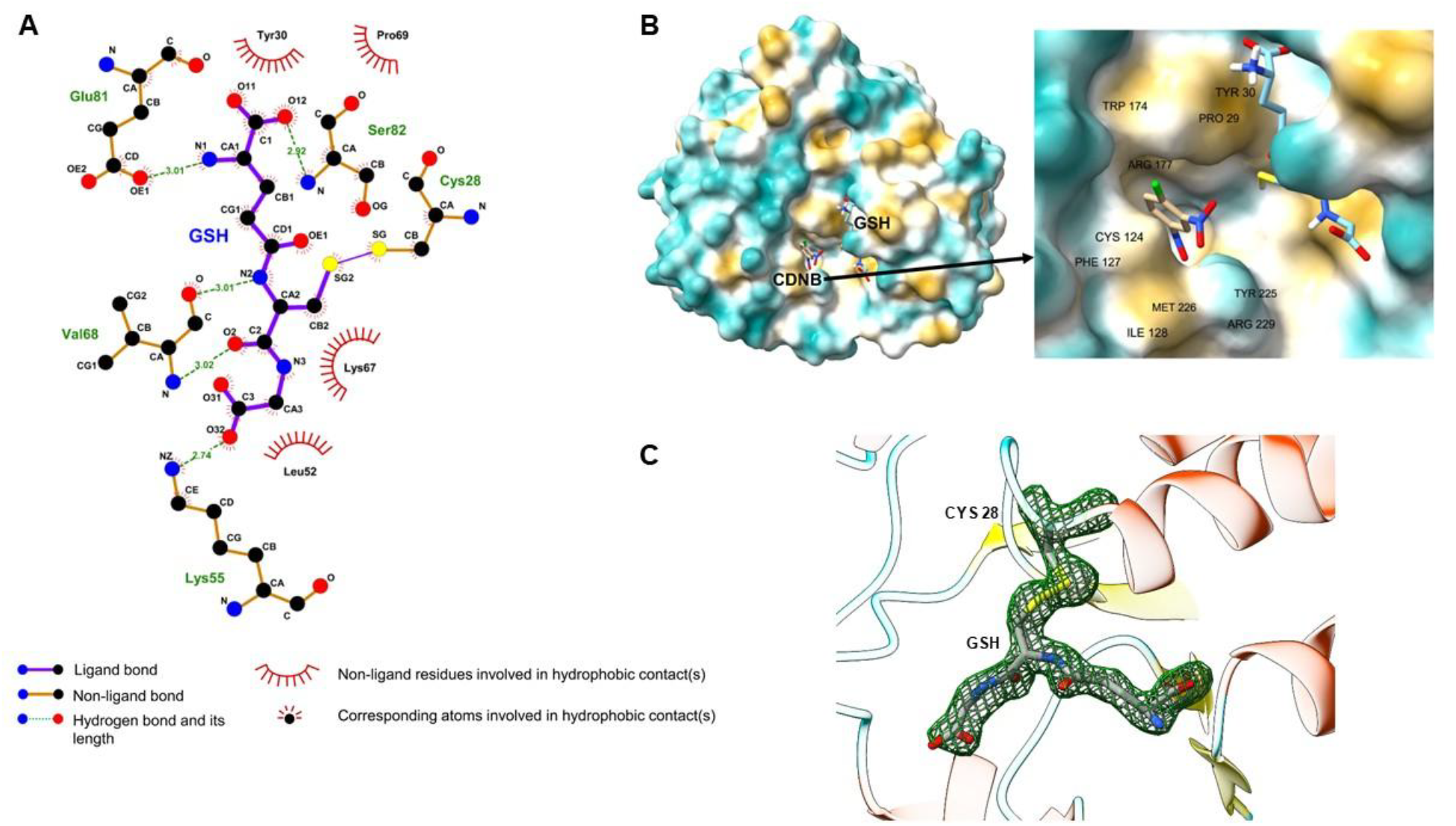
Protein-ligand interactions of AmGSTO1. (A) AmGSTO1-GSH interactions are represented graphically using LigPlot^94^. AmGSTO1 active site amino acid residues at G-site that bind GSH are shown. (B) AmGSTO1 active site amino acid residues at H-site that are in close contact with the docked ligand CDNB. (C) Ribbon diagram of AmGSTO1 zoomed in on G-site to display electron density for CYS28-GSH mixed disulfide. Polder map (mFo-DFc_polder) is displayed to show density within 3.5 Å of GSH and contoured ±3 σ.

### Enzymatic Kinetic of Wild-type and Mutant AmGSTO1

The kinetic parameters *V*_max_, *K*_m_, *k*_cat_, and *k*_cat_/*K*_m_ of AmGSTO1 towards CDNB, GSH, PNA and DHA were determined. Parameters were calculated in GraphPad Prism 9.5.1 by fitting experimental data to the Michaelis-Menten model using nonlinear regression (Fig. 5A-C). For various concentrations of CDNB and PNA while keeping the GSH concentration constant, *V*_max_ values were 30.21 ± 1.08 μM/min and 57.45 ± 5.09 μM/min; *K*_m_ values were 0.58 ± 0.05 mM and 0.69 ± 0.14 mM; *k*_cat_ values were 4.50 ± 0.16 min^-1^ and 2.14 ± 0.19 min^-1^; and *k*_cat_/*K*_m_ values were 7.76 mM/min and 3.10 mM/min, respectively (Table 2). When varying the concentration of GSH while maintaining a constant concentration of CDNB, the *V*_max_, *K*_m_, *k*_cat_, and *k*_cat_/*K*_m_ were 13.27 ± 0.49 μM/min, 0.03 ± 0.01 mM, 1.98 ± 0.07 min^-1^, and 66.00 mM/min, respectively (Table 2). The DHA reductase activity was determined at 1 mM DHA and 1 mM GSH. The initial velocity was 205.6 ± 1.9 µM/min and specific activity was 0.257 ± 0.002 µmol/min/mg. AmGSTO1 showed no detectable enzymatic activity towards isothiocyanate, 4-hydroxynonenal, and trans-2-hexenal (T2H).

**Fig. 5.**
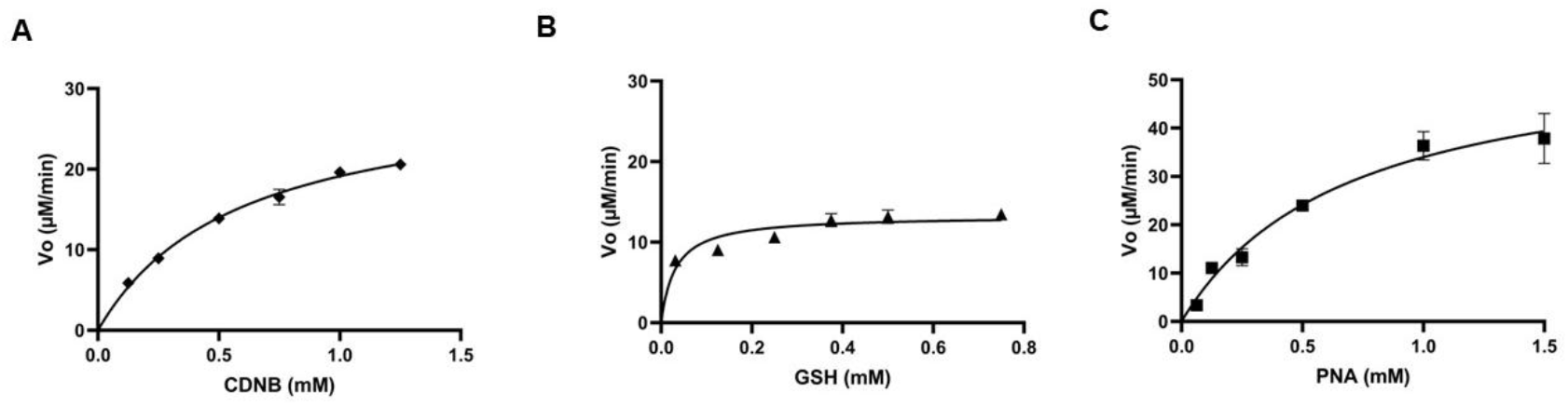
Plots of initial reaction velocity (V_o_) as a function of substrate concentration. (A) The CDNB conjugating activity at constant concentrations of GSH and various concentrations of CDNB from 0.125 to 1.25 mM. (B) The GSH-conjugation activity at various concentrations of GSH at 0.031 mM to 0.75 while holding the CDNB concentration constant. (C) The PNA-conjugation activity was tested using different concentrations of PNA ranging from 0.0625 to 1.5 mM, with the concentration of GSH held constant. *K*_m_ and *V*_max_ were calculated for each substrate using GraphPad Prism 9.5.1 by fitting experimental data to a nonlinear regression curve to obtain the Michaelis-Menten plot.

**Table 2.** Kinetic parameters for wild-type and mutant AmGSTO1 (WT – wild-type).

| H-site (Various concentrations of DHA, PNA, and CDNB with constant GSH concentration) |  |  |  |  |
| --- | --- | --- | --- | --- |
| | $V_{\max}$ ($\mu\text{M}/\text{min}$) | $K_m$ (mM) | $k_{\text{cat}}$($\text{min}^{-1}$) | $k_{\text{cat}}/K_m$ ($\text{min}^{-1}\cdot\text{mM}^{-1}$) |
| PNA | $57.45 \pm 5.09$ | $0.69 \pm 0.14$ | $2.14 \pm 0.19$ | 3.10 |
| CDNB |  |  |  |  |
| WT AmGSTO1 | $30.21 \pm 1.08$ | $0.58 \pm 0.05$ | $4.50 \pm 0.16$ | 7.76 |
| Y30A | $16.62 \pm 0.72$ | $0.21 \pm 0.03$ | $0.62 \pm 0.04$ | 2.95 |
| E81A | $31.81 \pm 1.60$ | $0.86 \pm 0.10$ | $1.19 \pm 0.06$ | 1.38 |
| F127A | $22.26 \pm 1.89$ | $0.64 \pm 0.15$ | $0.83 \pm 0.07$ | 1.30 |
| S82A | $29.02 \pm 2.06$ | $0.76 \pm 0.12$ | $1.08 \pm 0.08$ | 1.42 |
| W174A | $79.22 \pm 13.22$ | $2.28 \pm 0.68$ | $2.95 \pm 0.49$ | 1.29 |
| G-site (Various concentrations of GSH and constant CDNB) |  |  |  |  |
| | $V_{\max}$ ($\mu\text{M}/\text{min}$) | $K_m$ (mM) | $k_{\text{cat}}$($\text{min}^{-1}$) | $k_{\text{cat}}/K_m$ ($\text{min}^{-1}\cdot\text{mM}^{-1}$) |
| WT AmGSTO1 | $13.27 \pm 0.49$ | $0.03 \pm 0.01$ | $1.98 \pm 0.07$ | 66.00 |
| C28A | $263.85 \pm 21.98$ | $0.29 \pm 0.06$ | $9.84 \pm 0.82$ | 33.93 |

To assess the functional roles of active site residues (Cys28, Tyr30, Ser82, Phe127, Glu81, and Trp174) in AmGSTO1, each was substituted with alanine via site-directed mutagenesis, and kinetic parameters were measured. Replacement of Cys28 in the G-site (C28A) caused substantial changes in enzyme kinetics compared to the wild type. The *K*_m_ for GSH was 0.03 mM in the wild-type AmGSTO1 but increased 9.67-fold in the C28A mutant, suggesting decrease binding affinity (Table 2). The *k*_cat_ value for the C28A mutant increased by 4.97-fold compared to the wild type. Its catalytic efficiency (*k*_cat_/*K*_m_) decreased by 0.51-fold compared to wild type, suggesting impaired catalysis despite an increased turnover rate. Analysis of H-site mutants (Y30A, S82A, F127A, E81A, and W174A) revealed notable decreases in *k*_cat_ values. The greatest reduction in turnover was observed in the Y30A mutants (7.26-fold), followed by F127A (5.42-fold), S82A (4.17-fold), E81A (3.78-fold), and W174A mutant (1.52-fold) (Table 2). All G-site and H-site mutants showed reduced catalytic efficiency (*k*_cat_/*k*_m_) relative to the wild-type enzyme, highlighting the importance of these residues for catalytic activity of AmGSTO1.

### AmGSTO1 Binding to Selected Agrochemicals and Lack of Metabolic Transformation

Nonlinear regression analysis of ANS-AmGSTO1 saturation binding using a one-site binding model in GraphPad Prism 9.5.1 resulted in a *K*_d_ value 50.41 ± 2.67 µM (Fig. 6A). Various ligands and controls (GSH, CDNB, and EA) were evaluated as competitors of ANS bound at H-site of AmGSTO1 (Table 3). As a negative control, GSH showed no inhibition, confirming its lack of interaction with the H-site (Fig. 6B). In contrast, the positive controls, CDNB and EA, inhibited ANS binding, with IC_50_ values of 0.86 ± 0.02and 1.70 ± 0.07mM and had corresponding apparent *K*_i_ values of 434.34 and 858.59 µM, respectively (Fig. 6B, Table 3). Among 38 ligands screened, six compounds inhibited ANS binding by competing for the AmGSTO1 H-site (Fig. 6C). 2,4-D, fenoprop, 3,5,6-trichloro-2-pyridinol (TCP), 3-phenoxybenzaldehyde, tetramethrin, and nicotine showed IC_50_ values of 7.43 ± 1.07, 5.73 ± 0.83, 4.43 ± 0.48, 3.80 ± 0.30, 0.13 ± 0.01, and 4.30 ± 0.41mM, and corresponding apparent *K*_i_ values of 3752.53, 2893.94, 2237.37, 1919.19, 65.66, and 2171.72 µM, respectively (Fig. 6C, Table 3). HPLC MS/MS was conducted to evaluate the potential metabolism of fenoprop, TCP, and 2,4-D by AmGSTO1. Incubations with active and heat-inactivated GST showed no significant differences (*p*>0.05) (Fig. 7) indicating that AmGSTO1 is unlikely to metabolize these compounds directly.

**Fig. 6.**
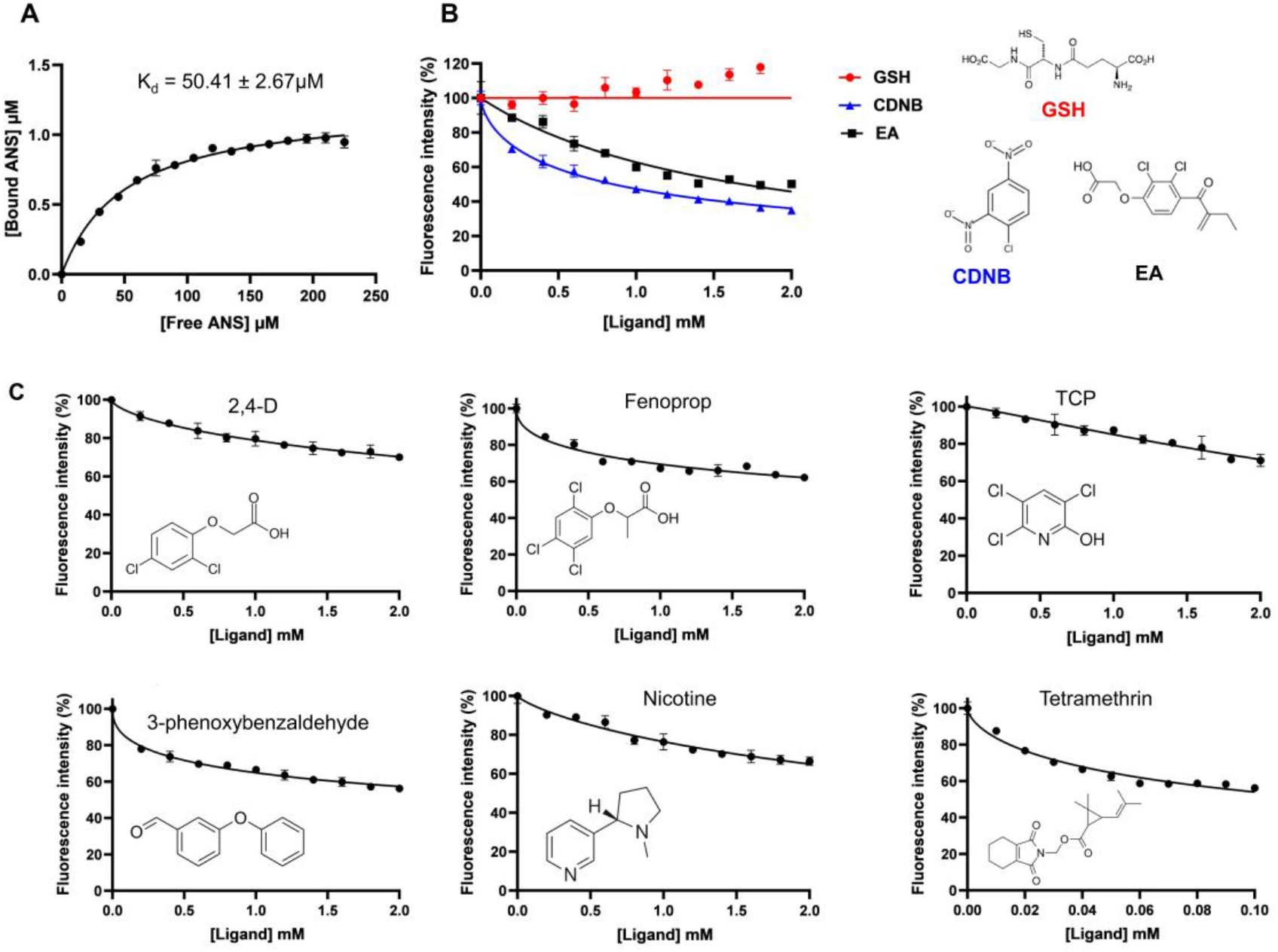
Fluorescence binding assay for AmGSTO1. (A) AmGSTO1 binding towards the fluorescent reporter 8-Anilinonaphthalene-1-sulfonic acid (ANS). Samples containing 2 µM/mL of AmGSTO1 were mixed with various concentrations of ANS in a buffer solution composed of 20 mM potassium phosphate, 150 mM NaCl, 1 mM EDTA, and 1 mM TCEP at pH 6. Fluorescence intensity was recorded using excitation and emission filters of 380 nm/20 nm and 485 nm/20 nm, respectively. Data analysis was performed using GraphPad Prism software with a one-site-specific binding model. (B) The displacement of ANS bound to AmGSTO1 was investigated using positive controls (CDNB and EA) and negative control (GSH). (C) The displacement of ANS bound to AmGSTO1 was examined using competitive ligands including 2,4-D, fenoprop, TCP, 3-phenoxybenzaldehyde, nicotine, and tetramethrin.

**Table 3.**
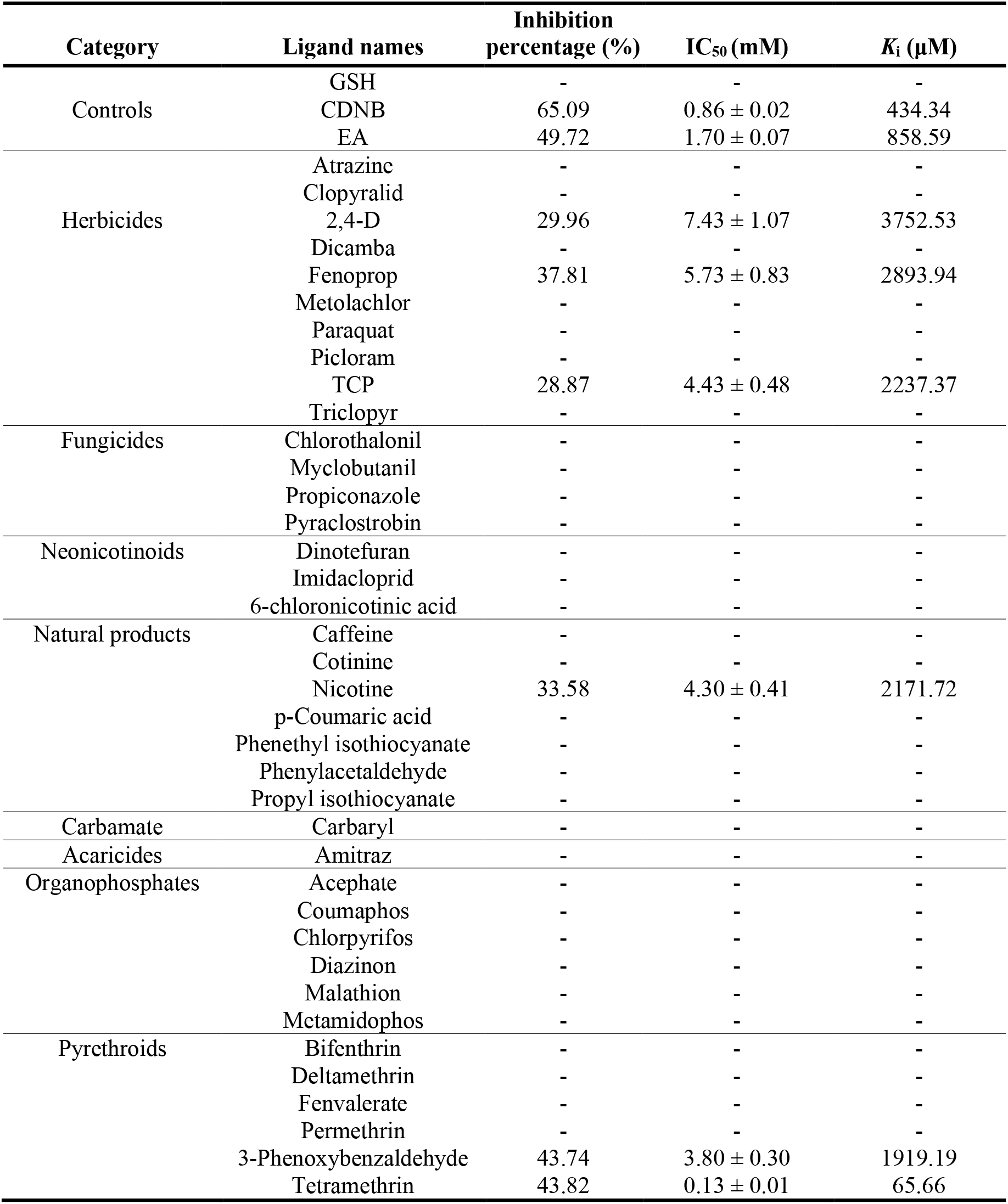
Competitive binding parameters of various ligands to AmGSTO1.

**Fig. 7.**
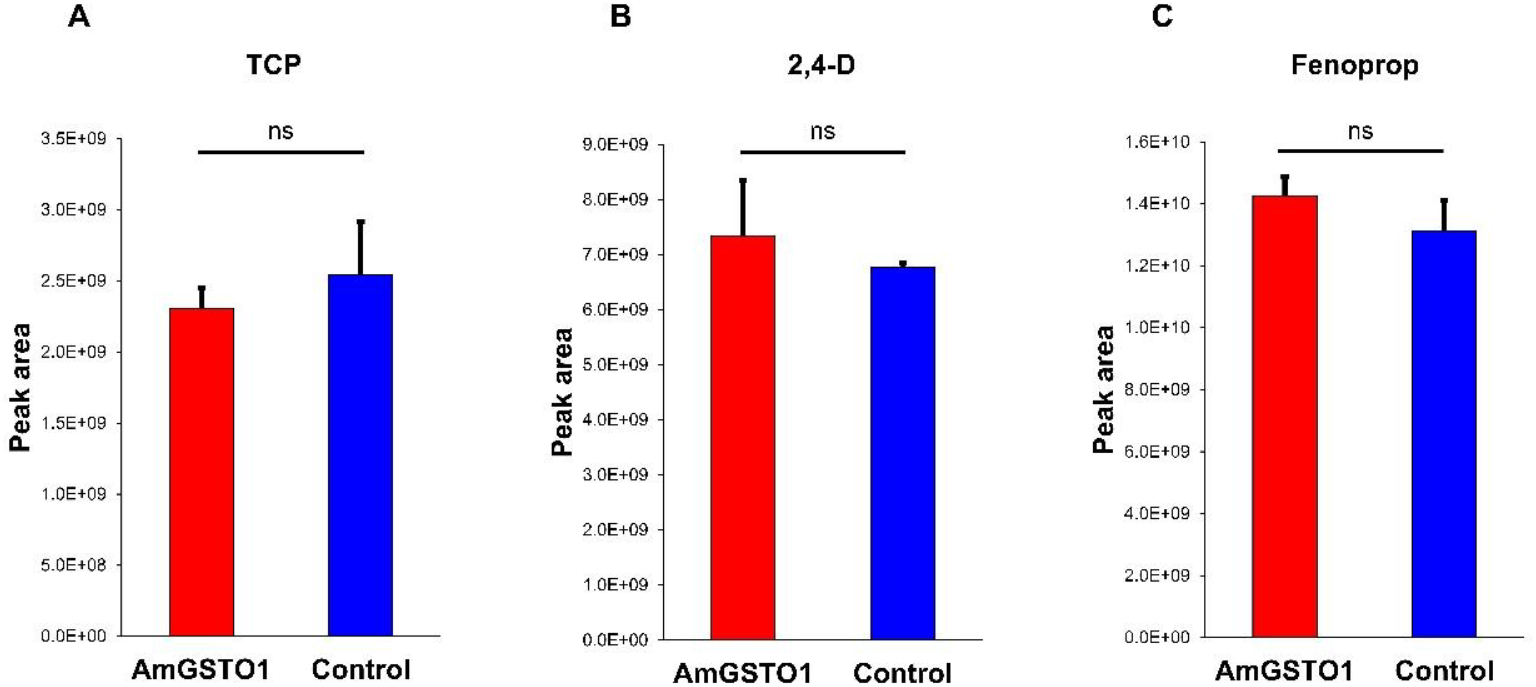
HPLC analysis and metabolic assay. The incubation of TCP and GSH with (A) heat-inactivated recombinant AmGSTO1 and (B) with functional AmGSTO1. (C) Bar diagram showing the amount of chemical substrates identified in the reaction mixture (left: TCP, middle: 2,4-D, and right: Fenoprop) after incubation with AmGSTO1 (treatment) and heat-inactivated AmGSTO1 (control).

### Antioxidative Properties of AmGSTO1

Disc diffusion and bacterial survival assays were used to evaluate the antioxidative role of AmGSTO1 against oxidative stress. CHP, H_2_O_2_, and paraquat were used as oxidative stress inducers ^60^. Disc diffusion assays revealed significant differences between the pET-9BC vector (control) and the recombinant AmGSTO1-pET9BC strains at all CHP (*p*<0.001; df = 9; F = 396.54) and H_2_O_2_ concentrations (*p*<0.001; df = 9; F = 295.15). After exposure to the CHP or H_2_O_2_, halo zone diameters were significantly smaller in LB agar plates containing AmGSTO1-expressing bacteria than in control plates (Fig. 8A-C, Fig. S2). Compared to the control, exposure to the highest concentrations of CHP and H_2_O_2_ (200 mM) resulted in reductions of 43.33% and 41.18% in average halo diameters, respectively. To complement the disc diffusion assays, bacterial assays were performed to quantitatively assess the antioxidative function of AmGSTO1. Survival assays revealed that the cells expressed AmGSTO1 exhibited significantly higher tolerance to H_2_O_2_ and paraquat than control cells (Fig. 9A-B). Increased OD_600nm_ in the treatment group compared with the control indicated enhanced bacterial survival. Colony-forming unit (CFU) counts following H_2_O_2_ or paraquat exposure showed survival rates of 90.4% and 83.3%, respectively in AmGSTO1-expressing cells, significantly higher than in controls (*p*<0.001) (Fig. 9C-F). Both disc diffusion and bacterial survival assays showed increased oxidative tolerance in AmGSTO1-expressing bacterial cells.

**Fig. 8.**
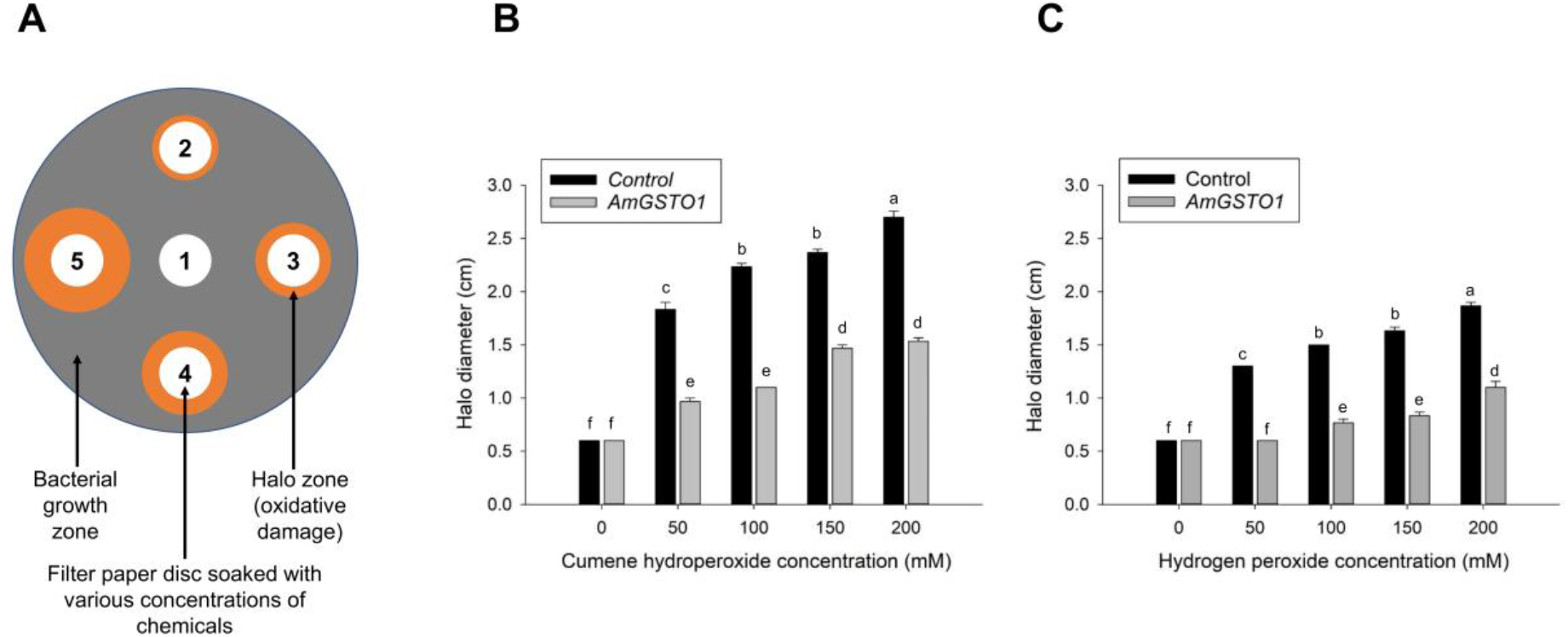
Disc diffusion assays in *E. coli* cells expressing AmGSTO1. (A) Cartoon image of filter paper disc soaked with varied concentrations of oxidative inducing chemical, *E. coli* cells growth in LB agar plate with antibiotics ampicillin (200 µg/ml) and chloramphenicol (30 µg/ml), and halo zone (clear area) created due to killing of cells by chemicals. (B) The halo zone developed under various concentrations of cumene hydroperoxide. (C) The halo zone developed under various concentrations of hydrogen peroxide. Different letters H_2_O_2_ Paraquat H_2_O_2_ Paraquat 90 indicate significant differences in halo zone formation between treatment and control as per one-way ANOVA with the Tukey HSD test.

**Fig. 9.**
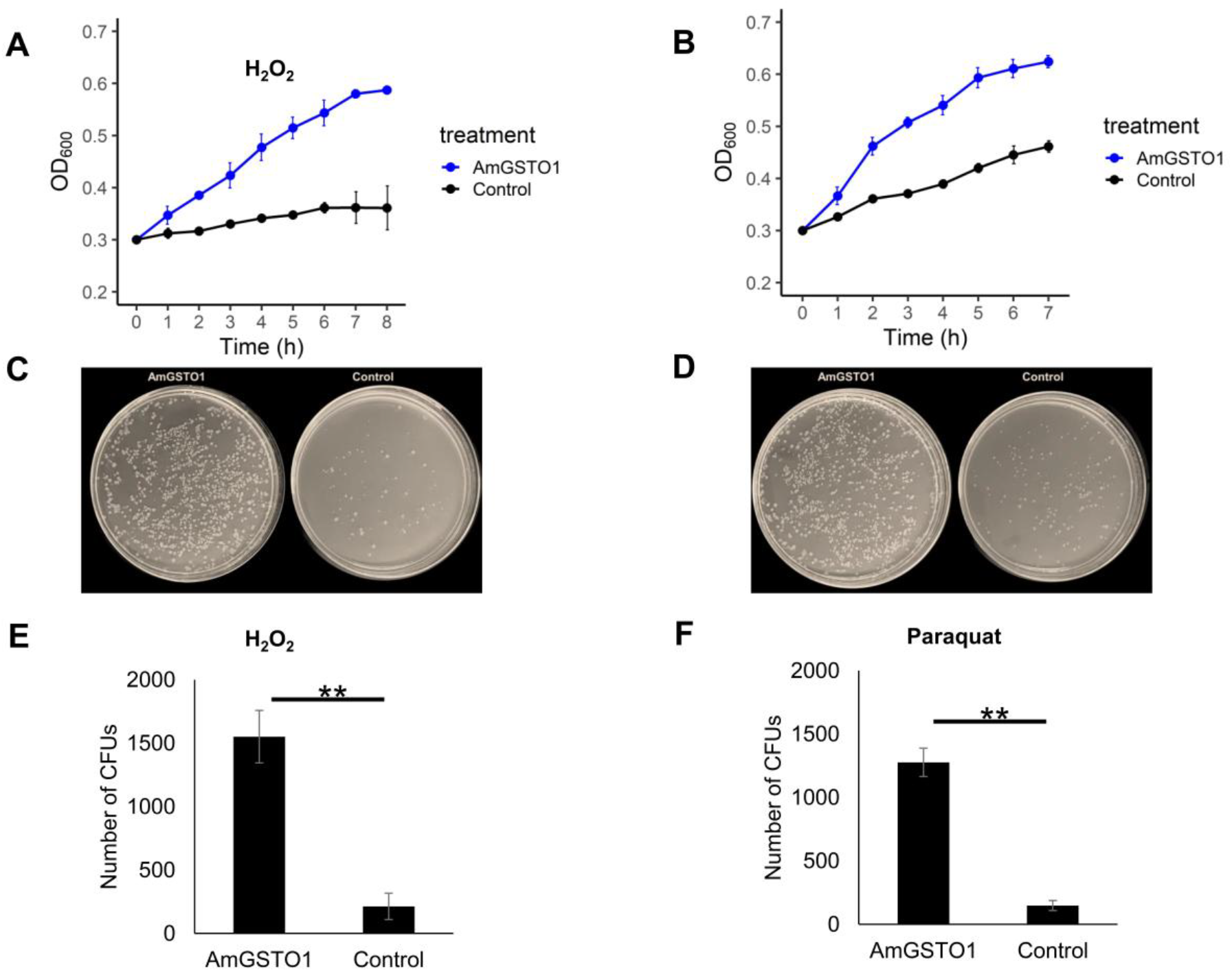
Bacterial survival assay in *E. coli* cells expressing AmGSTO1. *E. coli* cells consisting of recombinant AmGSTO1 and pET-9BC vector were grown in 2YT media containing H_2_O_2_ and paraquat (0.5 mM). The OD_600_ of *E. coli* was measured every hour following exposure to H_2_O_2_ (A) and paraquat (B) After 7-8 hours of incubation at 37 °C at 180 rpm, *E. coli* cells with recombinant AmGSTO1 and pET-9BC vector were smeared on agar plates and grown at 37 °C overnight (C and D). The number of bacteria colony-forming units (CFUs) on the agar plates was recorded under the microscope (E and F). ** Indicates a significant difference between the two groups as per Student’s t-test analysis (p < 0.001).

## Discussion

GSTs are central to detoxification processes and are thought to contribute to chemical adaptation in a variety of arthropod species ^20, 22, 25, 62-64^. In *Apis mellifera*, GSTs genes have been identified in different tissues and linked to diverse functions, including xenobiotic detoxification and olfaction ^17, 65-67^. Accumulating evidence from studies of Omega class GSTs in humans, microorganisms, and insects suggests that these enzymes contribute to oxidative stress mitigation ^59, 68-71^. Our study characterized the structure and function of the omega class GST, AmGSTO1, in honey bees. The findings presented here provide insight into potential mechanisms by which this enzyme could support honey bee adaptation to chemical stressors. In particular, AmGSTO1 may sequester pesticides and plant allelochemicals and mitigate oxidative stress, thereby protecting honey bee from diverse chemical stresses in their environment.

The co-crystal structure of AmGSTO1 bound to its co-substrate GSH reveals the amino acids that form the binding pocket. Our data showed that Cys28 in the G-site of AmGSTO1 is essential for GSH binding and activity with Cys28 forming a mixed disulfide bond with GSH in the co-crystal structure (Fig. 4A). Moreover, the Cys28A mutation significantly reduced catalytic efficiency (Table 2). This decrease in catalytic efficiency was primarily due to an increased K_m_ value in the C28A mutant, indicating a reduced GSH affinity after mutation, consistent with the Cys28-GSH mixed disulfide observed in the crystal structure. Omega GSTs are characterized by a conserved cysteine residue within the G-site, which facilitates the formation of a disulfide bond with GSH, in contrast with other GST classes that typically employ tyrosine or serine residues at the active site, highlighting a distinct catalytic mechanism from omega GSTs ^26, 59, 68^. AmGSTO1 exhibited significant DHA reductase activity, indicating a redox function as observed in other characterized omega GSTs ^55^. Additionally, AmGSTO1 displayed enzyme activity towards CDNB and PNA. In most omega GSTs, a proline residue commonly follows the conserved cysteine and is thought to stabilize the position of the cysteine thiol group, thereby supporting its chemical reactivity ^68, 72^. Consistent with this pattern, Pro29 occurs immediately after the Cys28 in AmGSTO1 (Fig. S3). Additionally, Arg33 and Leu38 were identified as conserved residues within the G-site of AmGSTO1 (Fig. 4C), similar to those reported for omega GSTs in *Homo sapiens* ^73^. This conservation suggests that these amino acids may be critical for omega GST functions across diverse species and may contribute to maintaining structure integrity. Adjacent to the G-site, a set of residues form a hydrophobic pocket, referred to as the H-site (Fig. 4B), which is thought to facilitate substrate binding and subsequent conjugation with GSH ^21, 74^. Alanine substitution of the H-site residues Tyr30, Ser82, Phe127, Glu81, and Trp174 led to reduced k_cat_ and k_cat_/K_m_ values, highlighting their importance for functional efficient catalysis. The decreases in turnover and catalytic efficiency Tyr30, Phe127 and Trp174 to alanine mutations point to the role of aromatic residues in defining the hydrophobic and productive geometry of the H-site for electrophilic aromatic substrates (Table 2).

Previous studies have employed competitive binding assays with hydrophobic fluorescent probes (e.g., ANS) to screen the substrate spectrum and binding capabilities of GSTs ^28, 39^. ANS binding studies showed that AmGSTO1 binds to the pesticides tetramethrin, 2,4-D, and fenoprop, to their metabolic byproducts: 3-phenoxybenzaldehyde and TCP, and to nicotine, a natural toxin found in *Nicotiana* pollen and nectar (Fig. 6C, Table 3). The pesticide tetramethrin displayed the strongest binding out of compounds tested, with an apparent *K*_i_ value of 64.21 µM. Pyrethroid insecticides are among the most frequently detected pesticide residues in beeswax, pollen, and honey bee-associated matrices in a previous study ^11^. Although pyrethroids are highly toxic to insects, including honey bees, their field application rates are typically low due to this potency and their reported repellent properties; however, increasing evidence indicates that even sublethal exposure can significantly impair bee physiology and behavior ^75-76^. In this study, selective binding was observed: tetramethrin and the deltamethrin metabolite interacted with AmGSTO1, whereas parent compounds such as deltamethrin and permethrin did not (Fig. 6C, Table 3). This suggests a degree of substrate specificity that may reflect differences in molecular size, polarity, or metabolic transformation. This pattern is consistent with the proposed role of omega-class GSTs in cellular defense processes, including antioxidant functions, rather than broad-spectrum xenobiotic conjugation ^71, 77-78^. In disc diffusion and bacterial survival assays, heterologous expression of AmGSTO1 increased tolerance to oxidative stress induced by common reactive oxygen species (ROS) generators, including paraquat, cumene hydroperoxide (CHP), and H_2_O_2_. Cells expressing AmGSTO1 exhibited improved growth and survival compared to control cells, suggesting that AmGSTO1 can enhance cellular resistance to oxidative stress in a bacterial model system (Figs. 8 and 9). Although demonstrated in a heterologous system, these findings are consistent with a potential antioxidant function of AmGSTO1 in honey bees under chemical stress conditions ^77^.

Besides insecticides, we also tested 10 commonly used herbicides and 4 fungicides (Table 3). Among them, two herbicides 2,4-D and fenoprop displayed 29.95% and 37.81% inhibition of ANS fluorescence, respectively, indicating their ability to interact with the hydrophobic binding site of AmGSTO1. TCP, a metabolite of herbicide triclopyr and a hydrolysis product of the organophosphate chlorpyrifos, also showed the ability to bind to AmGSTO1, exhibiting 28.85% inhibition of ANS fluorescence (Fig. 6C). Herbicides are generally considered to pose low acute toxicity to pollinators ^13^. However, exposure risk may increase substantially during the flowering period of crops or weeds attractive to bees. Elevated or repeated exposure has been reported to adversely affect bee health through mechanisms such as alterations in gut symbiont composition and disruption of colony-level processes including collective thermoregulation ^79-80^. When combined, multiple pesticides, herbicides and insecticides can produce synergistic or active effects that amplify toxicity and interfere with detoxification pathways, as reported in previous studies^81-84^. In our previous study, we found that AmGSTD1 exhibits binding affinity with 2,4-D, triclopyr, and TCP, as well as some insecticides and fungicides ^39^. Together with the present findings for AmGSTO1, these results suggest that honey bee GSTs contribute to chemical tolerance by interacting with a broad range of agrochemicals. However, competitive binding alone does not necessarily indicate metabolic activity. Although classical GST activity involves glutathione conjugation, there is precedent for GSTs interacting with xenobiotics in ways that do not result in direct metabolism. For example, in the fruit pest *Cydia pomonella (L*.), GSTs including omega-class family members were reported to bind the pyrethroid lambda-cyhalothrin without observable metabolic products, supporting the idea of sequestration in insecticide resistance ^85^. Similarly, an omega-class GST in *Anopheles cracens* Weidemann was shown to bind the organophosphate insecticide temephos without detectable glutathione conjugation, indicating that these enzymes can interact with xenobiotics through mechanisms other than classical catalysis ^86^. Catalyzed reactions indicate metabolic activity, in which enzymatic transformation of the ligand yields products that are often less toxic and more water-soluble, thereby facilitating excretion ^87-89^. Our binding assay, *in vitro* enzymatic assay coupled with HPLC, suggests that AmGSTO1 is likely involved in the sequestration of fenoprop, 2,4-D, and TCP but does not directly metabolize these chemicals (Fig. 7). GSTs binding to pesticides using a sequestering mechanism represents a passive mode of detoxification, which has been proposed in previous studies ^23, 85, 90^. From an evolutionary perspective, a sequestration-based mechanism may represent an advantageous strategy for coping with chronic, low-dose chemical exposure. Unlike catalytic detoxification, which requires precise substrate recognition and can generate reactive intermediates, ligand sequestration provides a passive and energetically efficient means of reducing the bioavailability of xenobiotics. For generalist pollinators such as honey bee, which encounter a chemically diverse landscape of plant allelochemicals and anthropogenic agrochemicals, selective pressure may favor detoxification enzymes with broad binding capacity rather than narrow substrate specificity. The ability of AmGSTO1 to bind multiple herbicides and pesticide metabolites without catalyzing their transformation is consistent with this model and aligns with the known functional characteristics of omega-class GSTs, which are frequently associated with redox homeostasis and stress tolerance rather than extensive xenobiotic conjugation ^91^. Together, these features suggest that sequestration-based interactions mediated by GSTs may have been evolutionarily favored as a flexible and protective response to environmental chemical stress.

Interestingly, nicotine, an alkaloid naturally occurring in *Nicotiana* nectar and pollen as well as in botanical insecticides, showed binding affinity towards AmGSTO1, exhibiting 33.57% inhibition of ANS fluorescence (Fig. 6C). This interaction may help to explain, at least in part, the tolerance to nicotine-containing host plants ^92-93^. As an agonist of nicotinic acetylcholine receptors (nAChRs), nicotine has been proposed to act as a reward component in nectar, potentially influencing foraging behavior through its involvement in neural signaling ^93^. Previous studies have reported upregulation of GSTs in honey bees following nicotine exposure, suggesting potential roles in Phase I and/or II detoxification processes ^65^. Taken together, the observed binding of both synthetic pesticides and plant allelochemicals in this study suggests that AmGSTO1 possesses a degree of structural flexibility that enables selective interactions with chemically diverse ligands.

In conclusion, this study characterized the tissue-specific expression patterns of *AmGSTO1* in nurse and forager honey bees and demonstrated its antioxidative capacity. Structural analyses identified key amino acid residues that are likely important for the catalytic and functional properties of AmGSTO1. Importantly, our findings suggest that AmGSTO1 may contribute to honey bee adaptation to agrochemicals and plant allelochemicals through a combination of selective ligand binding and mitigation of oxidative stress. Although AmGSTO1 did not directly metabolize several tested compounds, its binding capacity and antioxidant function support a role in cellular defense against chemical stress. Together, these results provide new insights into the structure–function relationships of omega-class GSTs in the honey bee and advance our understanding of how insects cope with diverse xenobiotic challenges.

## Acknowledgments

This work was supported by a faculty start-up fund from Pennsylvania State University, and the USDA National Institute of Food and Federal Appropriations under Hatch Project #PEN04938 and Accession #7006507 (F.Z.). T.W.M. was supported by USDA NIFA postdoctoral fellowship, grant #2020-67034-31780/project accession#1022959 (2020-2022) and USDA NIFA Hatch Project #PEN04897 and Accession#7005652 (T.W.M.). We are grateful to Avehi Singh, Kate Anton, and Margarita López-Uribe (The Pennsylvania State University) for their help in sample collection. We thank Tatiana Laremore (Proteomics and Mass Spectrometry Core Facility, Penn State) for their technical assistance with HPLC MS/MS. The crystal diffraction data was collected on beam time award GUP-71203 from the Advanced Photon Source using the APS Structural Biology Center’s (SBC) beamline 19-ID, a U.S. Department of Energy (DOE) Office of Science user facility operated for the DOE Office of Science by Argonne National Laboratory under Contract No. DE-AC02-06CH11357. We thank Michelle Radford and Youngchang Kim for their technical assistance with remote data collection at APS SBC beamline 19-ID.

## Supplementary data

**Table S1**. Primers used for this study, including qRT-PCR and full-length primers sequence for *AmGSTO1* as well as forward and reverse primers for creating mutations via substitution in active sites of GST genes. Nucleotide sequence with small letter indicates the codon targeted for mutation.

**Table S2**. Protein sequences for GST genes used to construct maximum likelihood (ML) phylogenetic tree.

**Figure S1**. SDS-PAGE for the final purified protein samples of AmGSTO1.

**Figure S2**. Disc diffusion assay for AmGSTO1. The agar plates coated with bacterial cells expressing either AmGSTO1 or the pET-9BC vector displayed qualitative observations of distinct halo zones when exposed to different concentrations of oxidative inducers. A and B: *E. coli* expressing pET-9BC and AmGSTO1, respectively, following exposure to cumene hydroperoxide; C and D: *E. coli* expressing pET-9BC and AmGSTO1, respectively, exposed to hydrogen peroxide.

**Figure S3**. Multiple sequence alignment of omega-class GSTs in honey bee (*Apis mellifera*) (AmGSTO1), human (*Homo sapiens sapiens*) (HsGSTO1 and HsGSTO2), and silkworm (*Bombyx mori*) (BmGSTO). Amino acids highlighted in green represent conserved amino acid residues at the G-site. HsGSTO1 (PDB: 1EEM_A), HsGSTO2 (PDB:3Q18_A), BmGSTO (PDB: 3WD6).

