## Supplementary Figures 1-3 for "An omega glutathione S-transferase in *Apis mellifera* contributes to chemical adaptation through pesticide sequestration and antioxidant defense"

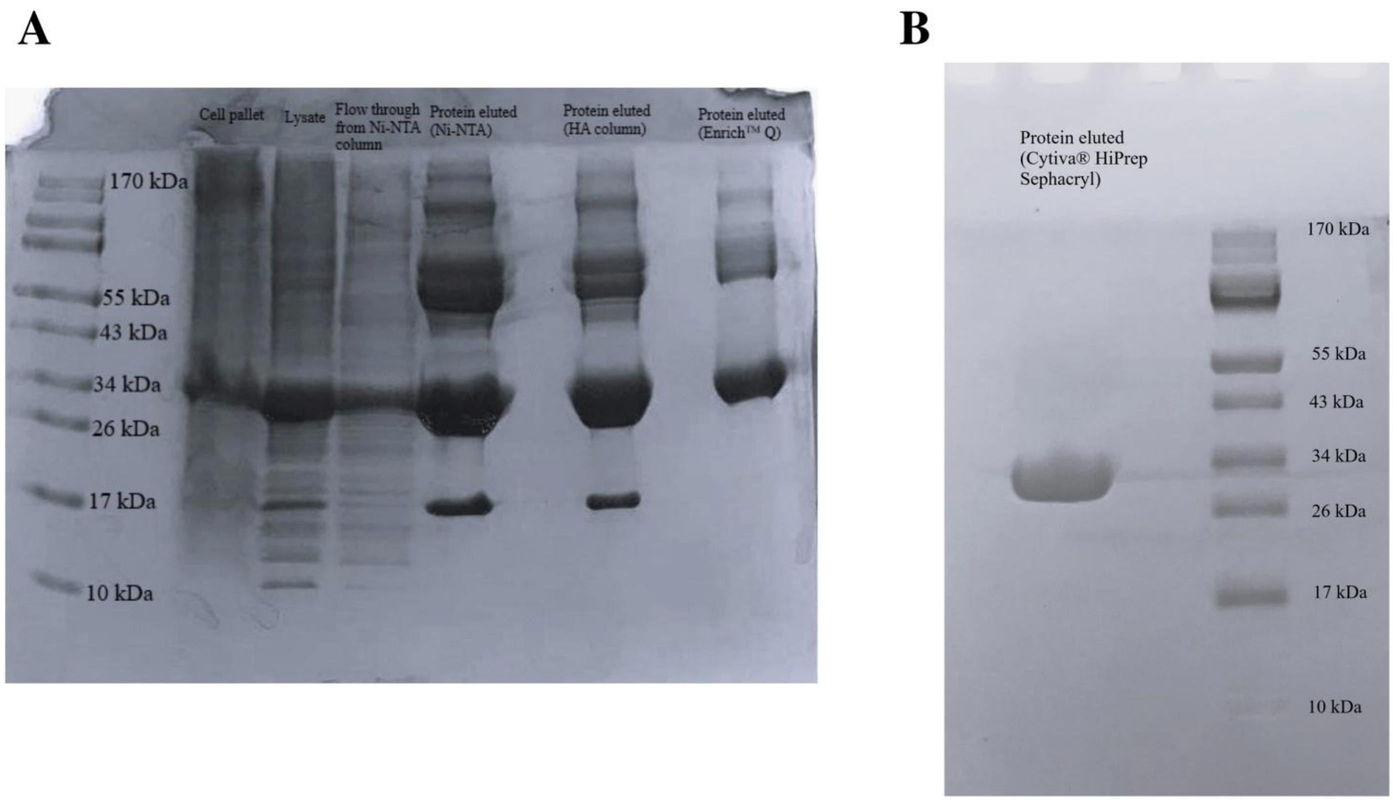


**Fig. S1.** SDS-PAGE for the final purified protein samples of AmGSTO1.

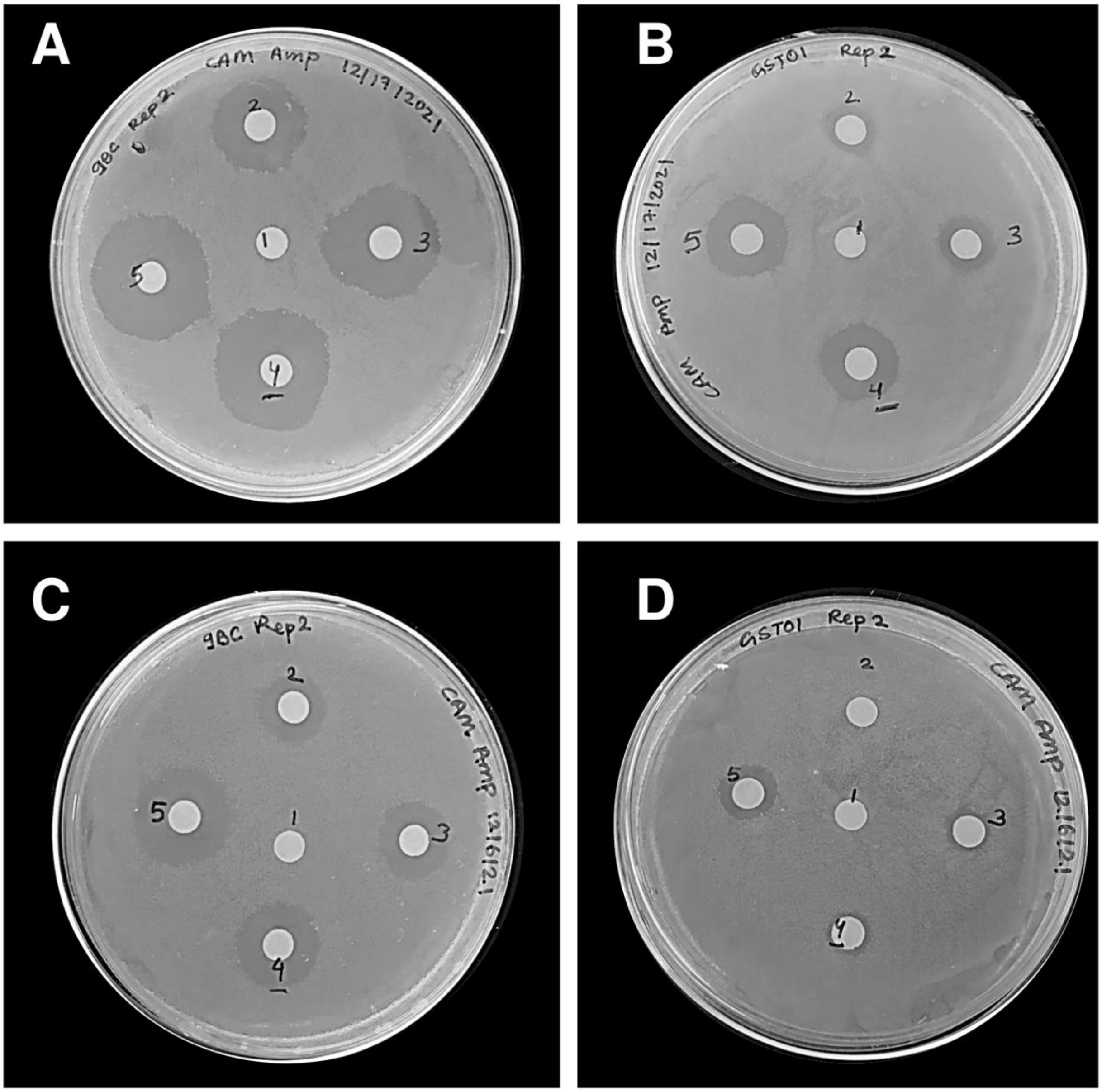


**Fig. S2.** Disc diffusion assay for AmGSTO1. The agar plates coated with bacterial cells expressing either AmGSTO1 or the pET-9BC vector displayed qualitative observations of distinct halo zones when exposed to different concentrations of oxidative inducers. A and B: *E. coli* expressing pET-9BC and AmGSTO1, respectively, following exposure to cumene hydroperoxide; C and D: *E. coli* expressing pET-9BC and AmGSTO1, respectively, exposed to hydrogen peroxide.

```
AmGSTO1      -----------MSSKHLTIGSVAP-PIVPGKIRLYSMRFCPYAQRIHLVLDAKHIPHDVV 48
BmGSTO       MSAIKDSRNINFNTKHLRKG--DPLPPFNGKLRVYNMRYCPYAQRTILALNAKQIDYEVV 58
HsGSTO1      MS--------GESARSLGKGSAPPGPVPEGSIRIYSMRFCPFAERTRLVLKAKGIRHEVI 52
HsGSTO2      MS--------GDATRTLGKGSQPPGPVPEGLIRIYSMRFCPYSHRTRLVLKAKDIRHEVV 52
```

**Fig. S3.** Multiple sequence alignment of omega-class GSTs in honey bee (*Apis mellifera*) (AmGSTO1), human (*Homo sapiens sapiens*) (HsGSTO1 and HsGSTO2), and silkworm (*Bombyx mori*) (BmGSTO). Amino acids highlighted in green represent conserved amino acid residues at the G-site. HsGSTO1 (PDB: 1EEM_A), HsGSTO2 (PDB:3Q18_A), BmGSTO (PDB: 3WD6).
